# If memory serves: Multisensory male display improves female memory during mate sampling

**DOI:** 10.1101/2025.06.25.660905

**Authors:** Olivia-Rose M. Hamilton, Logan S. James, Rachel A Page, Michael J. Ryan, Ryan C. Taylor, Kimberly L. Hunter

**Affiliations:** Department of Biological Sciences, Salisbury University, Salisbury, MD 21801, USA; Department of Biology, Wake Forest University, Winston-Salem, NC 27109, USA; Department of Integrative Biology, University of Texas, Austin, TX 78712, USA; Smithsonian Tropical Research Institute, Balboa, Ancón, Panamá

## Abstract

Across many taxa, males gather in leks to perform multisensory courtship displays for females. At a lek, changes in the sensory scene over the course of mate evaluation are inevitable. This dynamic nature makes a female’s ability to recall the location of individual signalers an important component of female mate choice. It is hypothesized that complex (especially multimodal) displays may improve a female’s ability to remember and thus discriminate amongst potential mates. To test this hypothesis, we presented female túngara frogs (*Physalaemus* (=*Engystomops*) *pustulosus*) with male calls (auditory) and robotic frogs (visual) that could later be obstructed from view via the lowering of blinds. Specifically, we asked if the visual component of a multimodal display improves the ability of a receiver to remember a signaler. While our initial experiment did not find memory instantiation in female frogs, subsequent experiments with a longer presentation time and a brief period of silence successfully instantiated memories in females. Moreover, females continued to demonstrate significant preferences for calls associated with the visual cue even after 25 s following the obstruction of the visual stimulus (robotic frog). Thus, we propose that the duration of presentation and/or the silent period impact memory capability. Silence is common in choruses, and our data suggest that complex, multisensory stimuli may have evolved to help females remember their preferred mate even with such fluctuations in signal production.

**LAY SUMMARY:** Multisensory signals function to stimulate one or more of a receiver’s senses, and in doing so, may make signalers more memorable to receivers. Multisensory signals are common in mating displays, including the advertisement of a male túngara frog: a call (acoustic) and a vocal sac inflation (visual). Here, we demonstrate that female túngara frogs were more likely to remember a multisensory male display over a unisensory display, but only after fluctuations in the signal’s intensity.

## INTRODUCTION

Animal communication signals constitute some of the most diverse traits observed in nature, many of which function in mate attraction (Bradbury & Vehrencamp 2011; Rosenthal 2017). Signaling reproductive availability can be achieved through a variety of behaviors and sensory modalities. The calls of crickets on a warm summer evening (Olvido & Wagner 2004), the low frequency “bellows” of a koala (Ellis et al. 2011), and the elaborate design of a male pufferfish’s nest (Kawase et al. 2013) are all produced in efforts to procure mates. Because females often decide the outcome of a reproductively motivated interaction, their decisions have critical fitness consequences (Andersson & Simmons 2006). For a signaler’s stimulus to be recognized, however, the nervous system of the receiver must integrate and analyze components of the signal and assign those components to their source (Bee & Miller 2016). Thus, the selective force of the receiver’s psychology, and the choice that is made as a result, is an important factor driving signal evolution (Guilford & Dawkins 1991).

Often, animals communicate using more than one sensory modality (Higham & Hebets 2013; Patricelli & Hebets 2016). During courtship displays, male butterflies, for example, use both chemical and visual cues (Costanzo & Monteiro 2007), while wolf spiders rely on visual and seismic signals (Kozak & Uetz 2019; McGinley et al. 2023). Male sagebrush lizards combine chemical and visual cues to both defend their territory and attract females (Thompson et al. 2008). Similarly, females prefer the visual display of a wingspread paired with an acoustic song over either stimulus alone in the brown-headed cowbird (Ronald et al. 2017). Bats, too, have been documented using multimodal courtship displays in which males use chemical, acoustic, and visual signals (Voigt et al. 2008). Simultaneous communication channels in different modalities are often presumed to provide more information, thereby improving a female’s ability to make critical mating decisions (Rosenthal 2017). It has therefore been predicted that the additional information in a multimodal display should aid the female’s ability to distinguish among individuals, particularly in noisy leks (Rowe 1999). Multisensory displays may also be more perceptible by a receiver, ultimately becoming more detectable in the environment. Utilizing a multimodal display may therefore confer benefits to males to make themselves more memorable when females are choosing among multiple options (Guilford & Dawkins 1991; Rowe 1999). Indeed, some studies have provided evidence for multicomponent displays improving a receiver’s memory both within and outside the context of mate choice (Rowe & Guilford 1999; Pardasani et al. 2021). In species with lekking behavior, the memorability of a signaler may be even more pertinent as females sample multiple males and make a choice within a relatively short time span (Backwell & Passmore 1996; Schwartz et al. 2004; Pauli & Lindström 2021). There are also periodic lapses in signal production (usually acoustic) (Greenfield 2015; Coss et al. 2020), so that during these quiet periods, females may have to rely on memory to make a choice or delay their decision. The “memory duration increase hypothesis” (Zhu et al. 2021) posits that eliciting increased memorability in receivers may contribute to the evolution of increased display complexity and multimodality.

Because females often assess multiple mates, learning and memory are important factors for female choice (Rowe 1999; Bateson & Healy 2005; Ryan et al. 2009), even on short time scales. For example, it is known that prior experience with conspecifics can influence mating decisions (Schlupp et al. 1994; Hebets 2003; Coleman et al. 2004; Cheetham et al. 2007; Bailey & Zuk 2009). Though several review papers have alluded to the possibility of learning and memory playing a role in the evolution of complex signals (Bateson & Healy 2005; Hebets & Papaj 2005; Bro-Jørgensen 2009; Higham & Hebets 2013; Lynch 2017; Ryan et al. 2019a; MacGillavry et al. 2023), few studies have experimentally tested this hypothesis. There are numerous studies across taxa that have investigated the role of spatial memory during mate searching in males (Jones et al. 2003; Goh & Morse 2010; Foley et al. 2015), but our knowledge about the general role of memory during mate searching in females is not as extensive (Healy & Hurly 2004). Specifically, one aspect of female mate choice that has received little attention is the role of multimodal cues in instantiating memory for signaler location. Females are likely to learn and retain information about a potential mate (location, size, signaling ability, etc.) so that they can recall it later. For instance, female house mice are able to recall the previous location of a male pheromone for 14 days (Roberts et al. 2012). For females that sample multiple males, especially over longer periods of time and across territories, it may be advantageous to remember information about each signaler to make a later choice, whether that be seconds from an initial contact or years.

The role of complex cues in memory during mate choice has primarily been performed using anurans. Anuran communication encompasses a diversity of display behaviors, with some species exhibiting visual, acoustic, chemical, and/or seismic signals and cues that are attended to by females (Gerhardt & Huber 2002; Höbel & Kolodziej 2013; Starnberger et al. 2014a; Robertson & Greene 2017; Caldwell et al. 2022; James et al. 2022). Despite this variety, acoustic signals remain the primary modality for most anuran communication, though these too can vary in their complexity (Reichert & Höbel 2018). In one of the studies investigating memory, Akre & Ryan (2010) found that female túngara frogs were able to recall the location of a more complex call for up to 45 s after cessation of the call. Yet, when presented with a simpler call, this effect was no longer exhibited, suggesting that females are able to maintain a working memory for the location of individual male callers. However, this study focused on the unimodal, acoustic component of the otherwise multimodal túngara frog mating display (Taylor et al. 2008). In a more recent study, Zhu et al. (2021) investigated the role of a multicomponent cue on the memory of the serrate-legged small treefrog (*Kurixalus odontotarsus*) and found that the multimodal display of a vocal sac inflation (via video playback) and an acoustic signal increased the active memory of females compared to a unimodal display. Because multimodal cues have been suggested to improve the memory of a receiver (Rowe 1999), these studies provide evidence that adding complexity to displays may improve the ability of a receiver to recall the last detected location of a preferred signaler. In the present study, we again rely on anuran communication to further our understanding of multimodal displays and memory.

Túngara frogs are a Neotropical species, with males gathering in nocturnal choruses to form leks (Ryan 1985). The acoustic stimulus of their mating display involves a simple “whine”, which may have up to seven additional “chucks” accompanying the whine (Ryan 1985). Females show a strong and consistent preference for a more complex call (whine plus chuck) (Ryan et al. 2019b). In addition to their acoustic stimulus, males produce cues in other modalities that can attract females. Consisting of an acoustic call, a visual cue, and a seismic component, the multimodal display of a male túngara frog is preferred over the acoustic call alone (Taylor et al. 2008; James et al. 2022). The male’s inflating vocal sac likely evolved to maximize calling efficiency, and its inflation secondarily became integrated into their mating display as the visual stimulus (Pauly et al. 2006; Taylor et al. 2008). The inflation of the vocal sac occurs synchronously with sound production during calling (Taylor et al. 2011). A túngara frog’s vocal sac is exceptionally large and, when inflated, is nearly the size of the male itself (Dudley & Rand 1991). In complex environments like the rainforests of Central America, the motion of a visual stimulus may be used by females to improve their ability to localize callers (Tan & Elgar 2021).

It is known that túngara frogs will stop calling in response to a predation threat (Dapper et al. 2011). Interruptions in the chorus are therefore common, with inter-chorus intervals having an average duration of 25 s (Akre & Ryan 2010). Because eavesdropping predators attend to the multimodal displays of males at a túngara frog lek (Halfwerk et al. 2014), females are vulnerable to predation while mate sampling (Bernal et al. 2007). It may be beneficial, then, for a female to evaluate males and make a choice quickly to mitigate predation risk. Experiments in both laboratory and natural settings seem to corroborate this (Rand et al. 1997). Female anurans tend to assess males and choose within a very short time period, with gray treefrogs (*Dryophytes (=Hyla) versicolor*) assessing potential mates for only about two minutes (Schwartz et al. 2004; Feagles & Höbel 2022). In dynamic lek environments, however, it is common for a male’s display to become interrupted or go silent during a female’s assessment. In that scenario, does she wait and restart her assessment once the chorus resumes, or does she rely on memory to influence her choice? As Akre & Ryan (2010) demonstrated, female túngara frogs are able to remember the location of a complex call for up to 45 s, spanning the average inter-chorus interval shown by the same study. Túngara frogs have also been shown to associate a visual cue with a place-learning task (Liu & Burmeister 2017; Burmeister 2022), suggesting they may depend more on landmark cues (i.e. a marker physically linked to a “goal”) than spatial cues (i.e. a marker that guides towards a “goal”) when remembering locations. It is possible, then, that the visual cue of an inflating vocal sac aids females in remembering a male’s call and, therefore, his location.

Here, we tested the hypothesis that the visual stimulus of an inflating male’s vocal sac instantiates female túngara frogs’ working memory for the stimulus, compared with a call only. In addition, we tested call playbacks with and without a silent period to better mimic natural settings and to understand how silence influences memory instantiation.

## METHODS

### General Experimental Procedures

All behavioral experiments were performed at facilities of the Smithsonian Tropical Research Institute in Gamboa, Panamá. We collected amplexed pairs of frogs from wild choruses between 1930 – 2300 h during the rainy season in 2021 and 2022. After transporting them to our lab for testing, frogs were stored in total darkness for at least one hour prior to trials to allow their eyes to dark-adapt. This dark adaption is necessary after collecting frogs using flashlights. For each trial, we separated a female from her mate and placed her under a visually and acoustically transparent mesh funnel in the center of a sound chamber (ETS-Lindgren, Austin, TX, USA).

Light inside the chamber was produced via a GE brand nightlight to mimic light conditions within the range of natural breeding conditions. The nightlight was green to the human observer, consisting of a broad spectral peak around 510 nm. This spectrum is within the range of what túngara frogs experience at breeding sites within forests and along forest edges (Taylor et al. 2008). We used tape to cover most of the nightlight surface, yielding a light-level of ~ 5.5×10^−10^ W/cm^2^. Light levels in the natural environment fluctuate widely depending on moon phase and cloud cover. Our experimental light levels were on the lower end of what the frogs experience in nature and within the range of the frogs’ scotopic visual sensitivity (Cummings et al. 2008; Leslie et al. 2021). The acoustic stimulus of each treatment was broadcast by a pair of Orb Audio speakers (Sherman Oaks, CA, USA; 1.2 – 18 kHz) that were placed 80 cm away from the release point of the female, forming an equilateral triangle (Fig. 1). For all experiments, the speakers antiphonally broadcast identical, synthetic calls (whine-chuck). A synthetic whine-chuck is, on average, as attractive as a natural call (Rand et al. 1992). We used Adobe Audition v22.2 to play call files. Once the female was placed inside the funnel, we presented the stimuli of each treatment (“presentation period”) for two to three minutes before the female was either 1) immediately released or 2) exposed to additional testing conditions and then subsequently released. After the female was released, we monitored and recorded her choice from outside the sound chamber using live infrared cameras. The video files were saved for an observer (blind to the treatments) to later rescore. We defined a choice as a female remaining within 8 cm of the base of a speaker (the “choice zone”) for at least 3 s. If a female failed to move from her initial “funnel zone” within 4 minutes or did not make a choice within 10 minutes, we removed her from the chamber. After a period of at least 15 minutes, the female was re-tested, and if she failed a second time, she was set aside and not tested again.

**Fig. 1.**
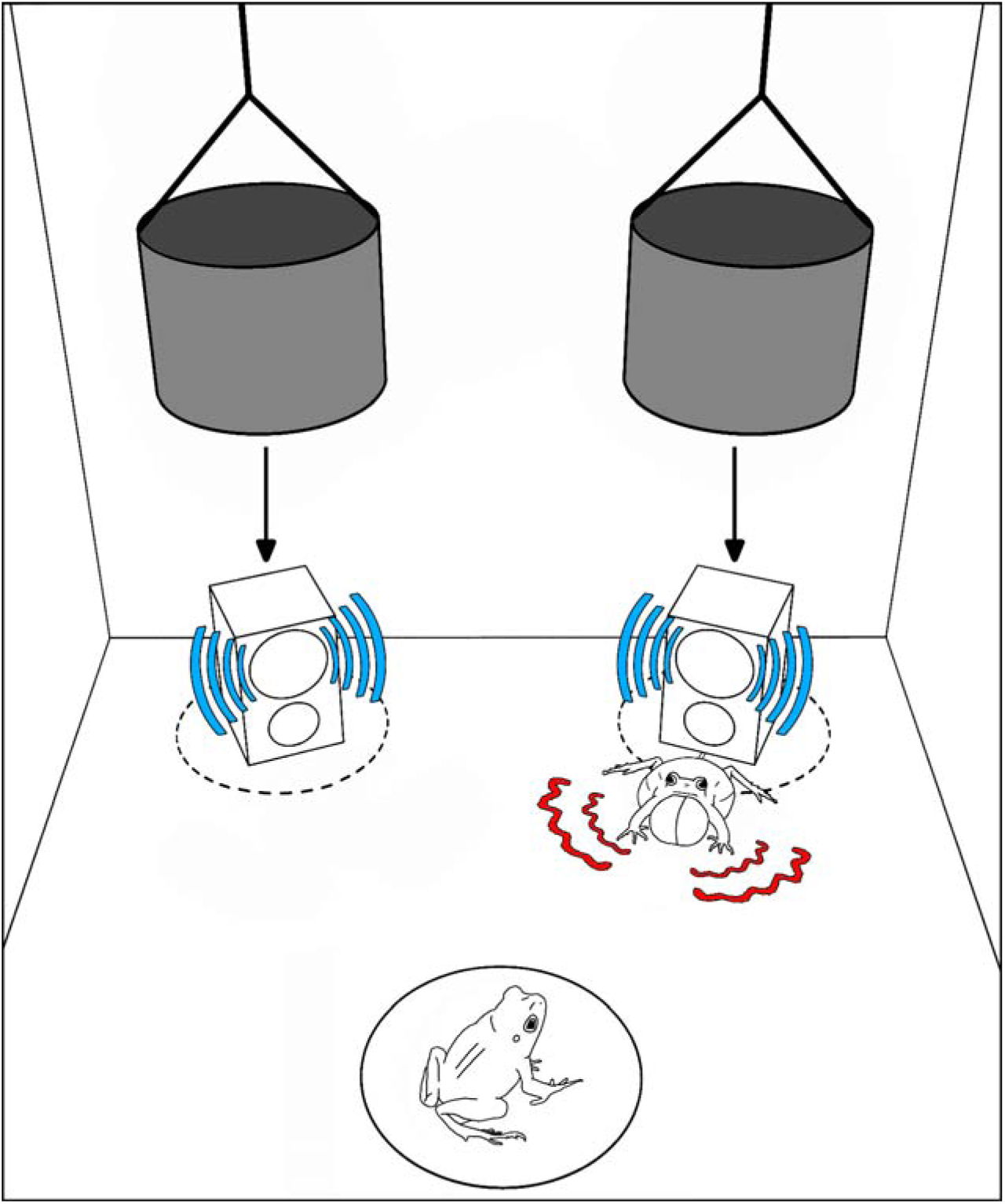
The internal arrangement of the sound chamber for memory phonotaxis experiments. (diagram not drawn to scale.) A female frog was placed in the center of the sound chamber in the funnel zone (black circle) and was presented with multimodal stimuli. The auditory stimulus (solid blue crescents) was broadcast by speakers while a robofrog simultaneously inflated a silicone vocal sac to serve as the visual stimulus (curved red crescents). After a presentation period, we gently lowered the blind cloths to obscure the visual stimulus from the view of the female, still held within the funnel zone.

We aimed to minimize our effect on the animals and their environment during testing. After completion of a trial, we reunited the female with her mate and later toe-clipped the pair, both to mark the frog (to avoid retesting the same individual) and to obtain a genetic sample. Toes were preserved in ethanol for future analyses. If females did not make a choice in an experiment, they were not toe-clipped. We followed regulations from the American Society of Ichthyologists and Herpetologists regarding toe-clipping procedures (Beaupre et al. 2004; Funk et al. 2005). Snout-vent-length and mass were also recorded for each frog. At the end of the night, all pairs were returned to their original collecting site and released. All frogs tested were unique individuals; females were not retested across treatments.

### Control: Robofrog Replication

Following Taylor et al. (2008), we replicated a multimodal stimulus for mate choice experiments using speakers and robotic frogs with artificial vocal sacs (Fig. 2a) inflated by an electromechanical pump (Fig. 2b). The robofrogs provided the visual component of the multimodal display while the call was produced by the speaker immediately behind the robofrog (Fig. 1). The pump was programmed to shunt air into tubing connected to the robofrog so that a silicone vocal sac inflated at a 19 kHz tone and deflated at a 16 kHz tone. These tones were placed at the respective beginning and end of the synthetic call in Adobe Audition. This ensured that the robofrogs inflated in time with the acoustic stimuli, as occurs in nature (Ryan 1985). These tones are imperceptible to frogs (Ryan et al. 1990). We calibrated the sound-pressure level (SPL) for both speakers at 82 dB SPL (re. 20 µPa, fast C-weighting) from the female’s position within the funnel. In this experiment, a robofrog was placed in front one of the speakers (right or left), and its position was switched between trials. During a two-minute presentation period, both speakers played a complex (whine-chuck) acoustic stimulus and the robofrog inflated synchronously with the speaker behind it. As both the acoustic and visual stimuli continued to play, the funnel was gently lifted from outside the chamber so that the female was free to make a choice. In addition to the definition of a choice used earlier, we also defined a choice in this experiment as the female approaching within 5 cm of the robofrog or touching it. As the robofrog was placed in front of a speaker, it extended the radius of the “choice zone” beyond that of a bare speaker. We note that the vocal sac movement (rather than the mere presence of the model frog) has been shown to be the critical aspect of the visual component, and must be closely timed with the acoustic call to induce a preference (Taylor et al. 2008; Taylor et al. 2011).

**Fig. 2.**
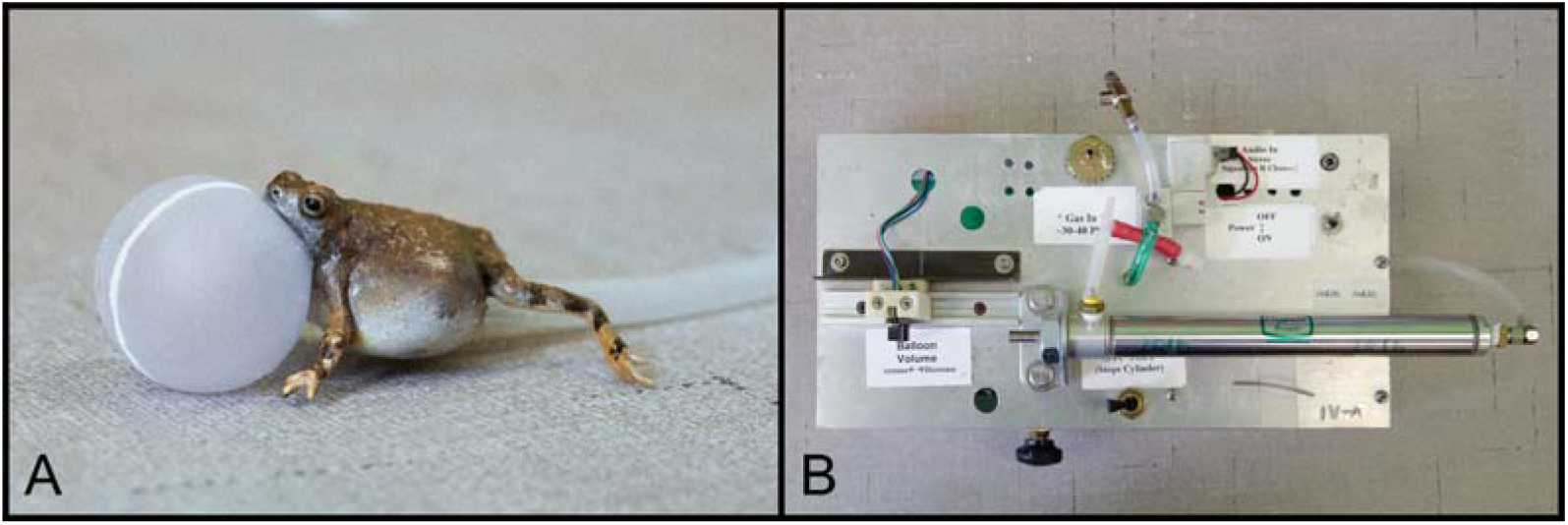
Robofrog setup. (A) Robotic, 3D-printed frog with inflated silicone vocal sac. (B) Robotic frog pneumatic controller. Compressed air drives the unit, and a solenoid valve controls the timing of inflation. During experiments, the pneumatic tube is hidden under a section of vinyl flooring that matches the floor of the test arena.

We tested a total of 46 individual females in this treatment. Thirty-two were tested in 2021. Data show that some climatic events may alter female túngara frog responses across years (Cronin et al. 2024). As such, we tested an additional 14 females in the 2022 robofrog control treatment to ensure that patterns of mate choice remained stable across the two years. All other experimental treatments were conducted in 2022.

### Memory Tests

For all subsequent treatments, we tested whether females employ their memory of the visual stimulus of a calling male (robofrog) that is suddenly obstructed from view when choosing between two sound sources. An opaque, black cloth served as a blind to block the visual cue from all angles (Fig. 1). The blind wrapped around both the speaker and the robofrog (if present) 360° to completely obscure the female’s ability to see the speaker/robofrog, irrespective of her position in the test chamber. We designed the blind to be as tall as the speakers (13 cm) so that the females could not see anything that was hidden behind them. The cloth was hand-sewn to a metal ring, which was then tied to monofilament fishing line. The fishing line was accessible to the outside of the chamber through a small port that also allowed access for speaker cables. With the blind surrounding the speakers, the amplitude at the female’s release point was 76 dB SPL; a decrease of 6 dB from a bare speaker without the blind. This amplitude is still within the range of what females experience in nature and has been used in previous studies in which females continue to exhibit robust preferences (Stange et al. 2016). In addition, both speakers were always enclosed within the blind, thus the drop in amplitude was identical for both speakers. During the presentation period (initial playback to a female), the blinds hung unmoving from the ceiling of the chamber. We then lowered the blinds to obscure the robofrog. As the blinds were lowered, an observer at the live camera feed monitored their rate and speed to ensure they were even and touched the floor of the chamber simultaneously. The lowering of the blinds took approximately 7 s. Because the original choice zone used for the speaker was engulfed by each blind, we defined a choice as a female approaching within approximately 8 cm of a blind and remaining there for at least 3 s. Despite the fact that the robofrog was occluded by the blind, we still halted the inflation and deflation of the vocal sac whenever the blinds were lowered.

#### Control: Blinds

We first confirmed that females would continue to demonstrate typical phonotaxis behavior despite the speakers being occluded by the blinds. For this control experiment (n=13), each speaker played an identical whine-chuck call for the entirety of the experiment. Our sample size for this control was smaller than other treatments, as we were limited by time during our field season. We did not utilize a robofrog as we were simply verifying that females would exhibit phonotaxis behavior and that there was no aversion or bias for one cloth over another. After the presentation period, the blinds were lowered, and once the blinds touched the floor, the female was immediately released from the funnel.

#### Memory: Hold and/or Silence

All memory tests began with a presentation period, during which the speakers antiphonally broadcast identical whine-chuck calls. The presentation period lasted 2 min for the non-silent treatments and 3 min for the silent treatments. The increase in time for the silent treatments was to ensure that a data collector could locate the monofilament lines in a dark laboratory and be prepared to lower the blinds in a consistent manner. Traditionally, a 3 min acclimation period has been utilized in túngara frog phonotaxis studies (Bosch et al. 2000; Marsh et al. 2000; Baugh & Ryan 2011). One robofrog inflated in time with the call of the speaker it was paired with and was visible to the female during this period. We switched the robofrog between speakers across trials. After the presentation period, the blinds were lowered and females were exposed to one of three conditions: a silent treatment, a hold treatment, or a combined silent & hold treatment (Fig. 3).

**Fig. 3.**
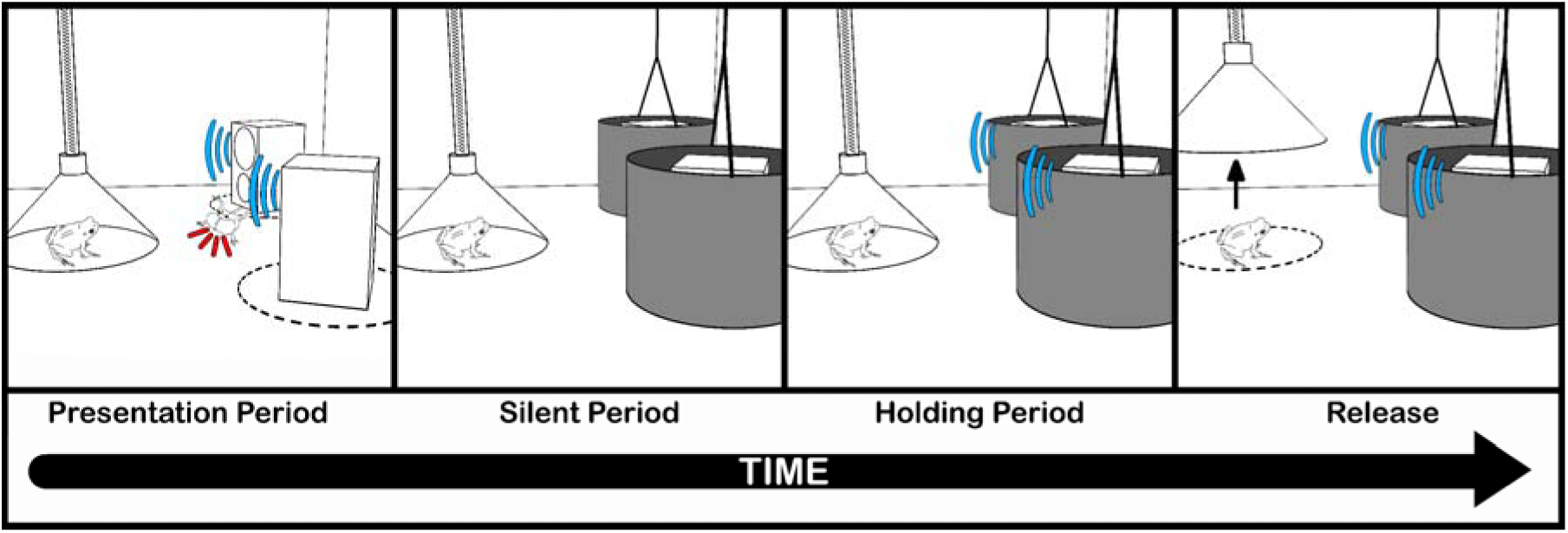
Periods of stimuli for female frogs in our experiments (image not drawn to scale). All experiments first involved a presentation period of 2 or 3 min where the female was held in the funnel and speakers antiphonally broadcast attractive (whine-chuck) calls (depicted by the blue crescents). We placed a robofrog (visual stimulus, depicted by the red lines) at one speaker that had a silicone vocal sac inflating in time with the call at that speaker (multimodal speaker). Immediately after the presentation period, the blinds were lowered and the females experienced a silent period, holding period, or both. The silent period of 5 s had no stimuli (acoustic or visual) presented to the female. For the holding period, the visual stimulus was no longer presented to the female, but the acoustic stimuli were broadcast. The holding period lasted for 20 s. Immediately following one or both of these periods, the acoustic stimuli either began or continued to play and the female was released from the funnel. From here, the female was free to make a choice between speakers.

For the hold treatment, as soon as the blinds touched the floor, females were retained under the funnel for 20 s but call playbacks from the speakers continued uninterrupted (n=38). After this 20 s, the female was released and allowed to make a choice (Fig. 3).

For the silent treatment, once the blinds touched the floor, all sound was muted from the speakers for 5 s (n=25). After this 5 s silent period, the female was released, and the audio (but not visual) playback began again and continued to play until the female made a choice (Fig. 3).

For our final treatment, we combined the silent and hold conditions in sequence (n=25). This treatment was the longest test of memory, with 25 s total time between the lowering of the blinds and the females’ release from the funnel. Females in this treatment first received the silent period. After the 5 s silence period, the funnel remained in place and the acoustic playback resumed. Females remained in the funnel for 20 s, as in the hold condition, before the funnel was raised (Fig. 3). In all memory treatments, the acoustic playback continued until the female made a choice or the 10-minute time limit was reached.

### Locomotor Behaviors

In addition to choice and time to choice (latency), we also analyzed the path females took to reach a speaker and the number of “stops” and “turns” they made on the way to a speaker. Using our recorded videos, we had an observer (unaware of the hypothesis) score choice and movement behaviors. We divided the floor of the test chamber into quadrants with the center converging on the funnel at the female’s release point (Fig. 4a). If the path the female took to reach a speaker remained within the same quadrant as the speaker, we scored that as a “direct” path. If, at any point, the female walked/hopped into a different quadrant than the speaker she ultimately chose, we scored that as an “indirect” path (Fig. 4a). We defined stops as any cessation of movement for >1 s. Turns were scored when a female changed the angle of the long-axis of her body 30° or more (Fig. 4b). Intuitively, frogs taking an indirect path are likely to exhibit higher numbers of stops and turns (versus the direct path). Frogs taking the direct path, however, still had sufficient space and time to express these behaviors. Locomotor behaviors have the potential to provide additional insights into the behavior of mate searching females. These behaviors are typically not scored in anuran phonotaxis tests (but see Feagles & Hobel 2022; 2023).

**Fig. 4.**
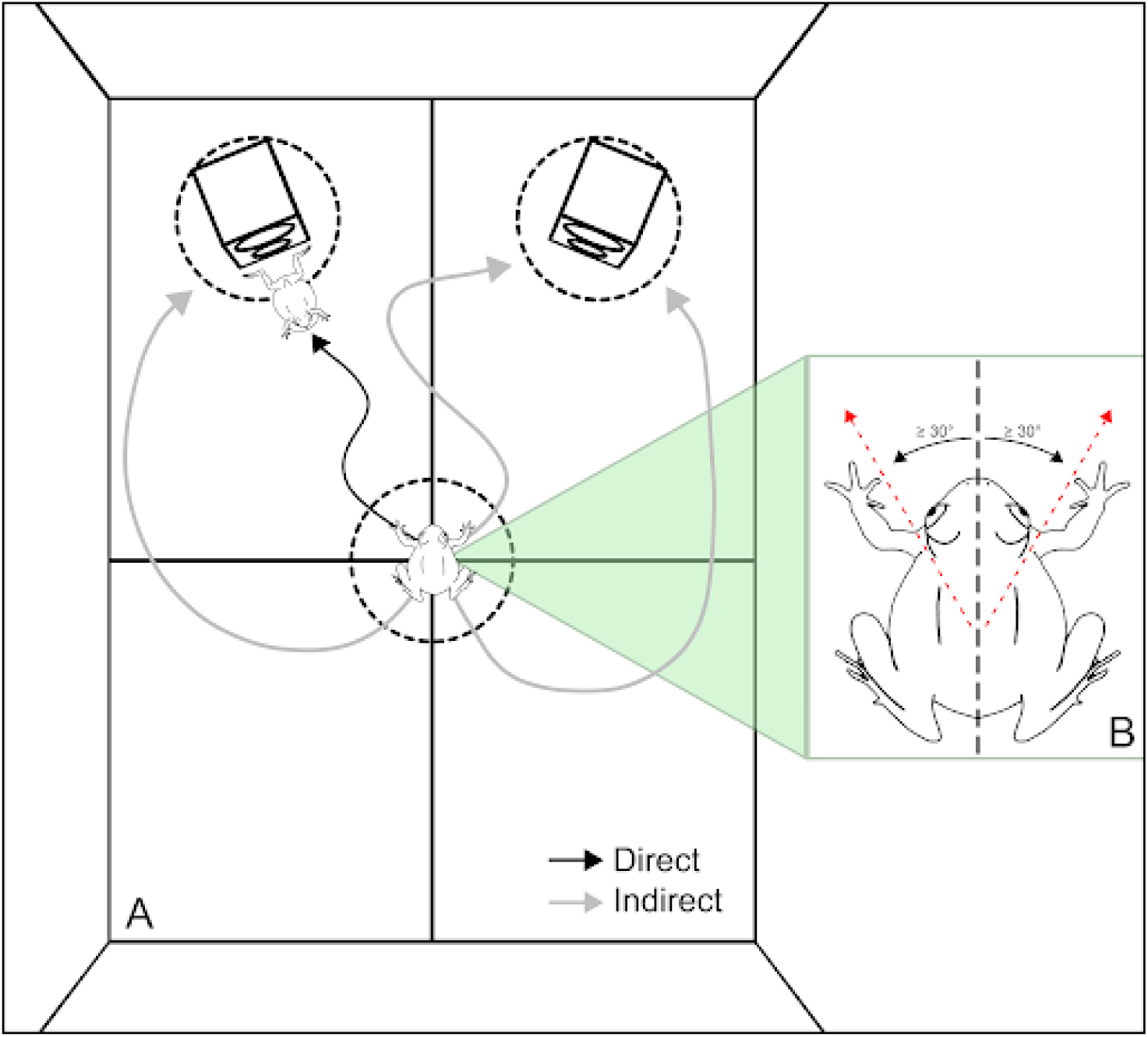
(A) View of the sound chamber from above, separated into four quadrants (figure not drawn to scale). If at any time during a trial a female left the quadrant containing the speaker she chose, we considered her path to be indirect regardless of the robofrog’s location. (Quadrant lines were used by a blind observer to visualize a female’s path; the lines were not present during phonotaxis experiments.) Possible paths out of the funnel zone are represented by either gray arrows (indirect path) or black arrows (direct path). (B) The dorsal view of a female túngara frog. When scoring videos, the observer quantified how many stops/turns a female would make. A movement was counted as a turn if the frog oriented her body 30° or more from the long-axis of her body.

### Ethics Note

Experimental procedures were conducted with approval from Salisbury University and the Smithsonian Tropical Research Institute (IACUC: SU-0052; SU-0052R and STRI 2018-0411-2021; SI-21012). The Ministry of the Environment of Panamá (MiAmbiente) approved our collection of animals and issued collecting permits for our team (ANAM: SE/A-39-2020). Procedures for marking animals were conducted in accordance with the American Society of Ichthyologists and Herpetologists.

## STATISTICAL ANALYSES

We first tested the hypothesis that the proportion of choices for each treatment were different from random chance using a binomial test. We report the mid-P-value to smooth large jumps in p-values, as often occurs with categorical data (Agresti 2001; Hwang & Yang 2001). Next, we used a Kruskal-Wallis test to examine how the number of stops and turns an individual made during a trial varied depending on whether she took a direct or indirect path to her choice. A binomial test was also used to compare the path of females (indirect vs direct) across treatments. We used an ANOVA to compare the mean latency to choose across treatments. To test if choice preferences were different across years of data collection, we performed a Fisher’s exact test (SISA Fisher exact test; Uitenbroek DG 1997). All other analyses were performed in R (R Core Team 2020). Graphs were generated in R (R Core Team 2020) using the packages: tidyverse (Wickham et al. 2019), ggplot2 (Wickham 2016), and ggpubr (Kassambara 2023). Figures were edited using Blender v4.2.0 and Adobe Photoshop v25.11.

## RESULTS

We tested female túngara frogs in several conditions to determine the role memory played when choosing between two speakers, where one speaker was formerly paired with a visual cue. Using a blind (opaque cloth), we occluded the visual stimulus (robofrog) from the view of the female in three different memory conditions.

First, we replicated the previous finding that females (*n* = 46) preferentially choose a speaker paired with a dynamic robofrog (multimodal display) over a bare speaker when both speakers played the same call (whine-chuck). Because we tested frogs in both 2021 and 2022 for this treatment, we wanted to ensure that their collection year did not influence preference. Though we know that preferences are stable and robust across years (Ryan et al. 2019b), a recent study demonstrated that drought can influence females’ preferences between field seasons (Cronin et al. 2024). Across the two years, the preference ratio was 22:10 in favor of the robofrog (*p* = 0.043), and 11:3, in favor of the robofrog (*p*= 0.046). A Fisher’s exact test showed that these ratios were not significantly different (Pearson’s χ^2^; *p* = 0.496), so we combined both years for the binomial test. Overall, females expressed a significant preference for the multimodal display over the unimodal acoustic call (binomial test: 33 robofrog:13 unimodal *p* = 0.0024; Fig. 5), similar to what we have shown elsewhere (Taylor et al. 2008; Stange et al. 2016).

**Fig. 5.**
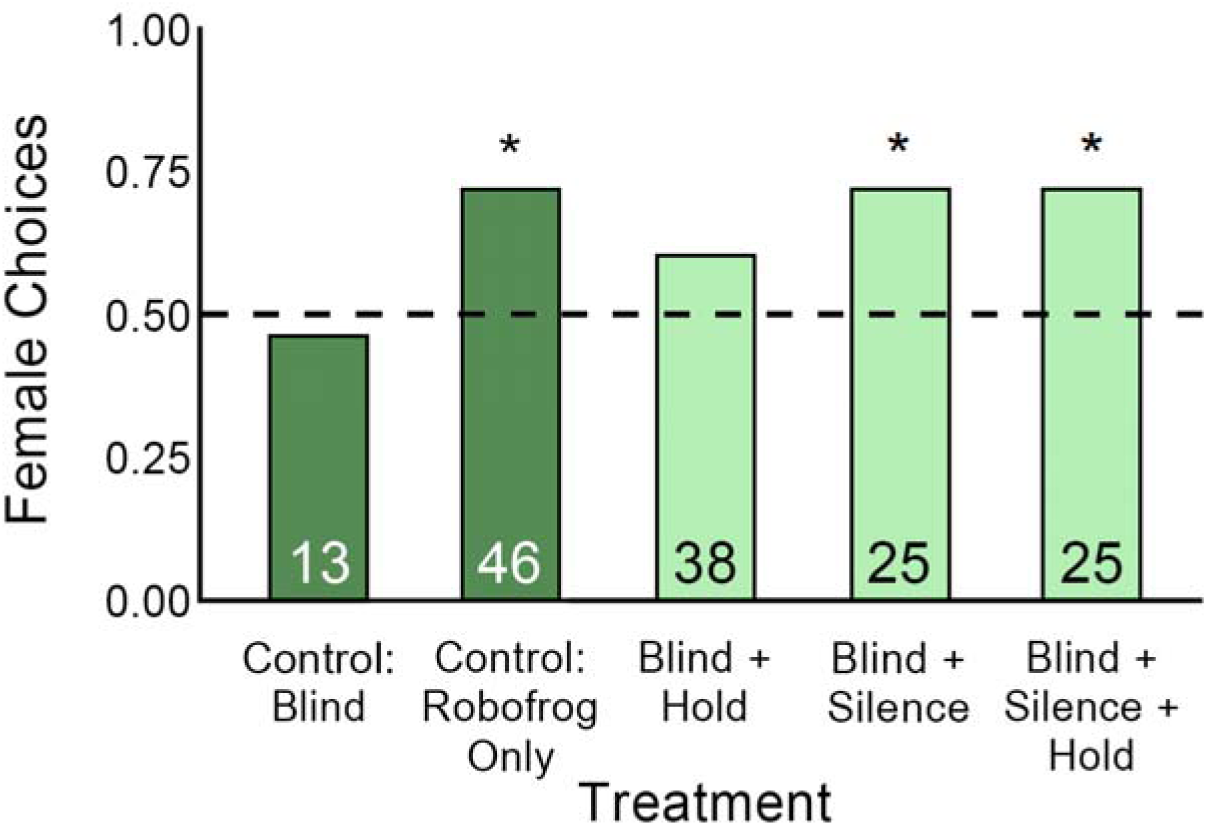
Proportion of female choice across treatments. Dotted line represents a random preference (50%). Asterisks (*) above bars indicate a significant difference from the expected preference of equal probability = 0.5. Numbers inside bars represent sample sizes for each respective treatment. The dark green bars (left) denote control treatments.

Next, we confirmed that the use of blinds had no impact on phonotaxis behavior. Despite the decrease in call amplitude after the blinds surrounded the speakers, females continued to demonstrate typical phonotaxis behaviors. Furthermore, there was no side bias for one cloth over the other (*n* = 13; binomial test: 6:7 ratio; *p* = 0.8953; Fig. 5).

We ran our primary experiments testing whether females prefer a speaker where they had previously been able to see the visual component of the robofrog. For all memory experiments, one speaker was paired with a dynamically inflating robofrog during the presentation period. At the end of the presentation period, we lowered the black cloths and halted the robofrog inflation, at which point females experienced one of three treatments: silence, hold, or a combined silence & hold.

Females in the hold treatment experienced a 2-minute presentation period and a 20 s holding period after the visual stimulus was hidden, but experienced no break in the auditory stimulus. Data for this treatment were collected in both 2021 and 2022. The ratios (robofrog : bare speaker) for the two years were 2:4 and 21:11. A Fisher’s exact test showed that these were not significantly different (Pearson’s χ^2^, *p* = 0.14). We combined both years for the binomial test. Overall, females did not approach the multimodal speaker significantly more often than random (binomial test: 23:15; *p* = 0.228; Fig. 5).

Females in the silent treatment (*n* = 25, all data collected in 2022) experienced a 3-minute presentation period and a 5 s silent period which began as soon as the visual stimulus was hidden. Subsequently, the females were released as the acoustic stimuli began playing again. Females chose the speaker that was formerly paired with the robofrog (multimodal speaker) significantly more often than the bare speaker (binomial test: 18:7; *p* = 0.0361; Fig. 5).

Our final group of females experienced a combined treatment (*n* = 25, all data collected in 2022). They first experienced a 3-minute presentation period, then the 5 s silent period, followed by a 20 s holding period where a female was held under the funnel while calls continued to play. After the holding period, the female was released and allowed to make a choice. Even after a total of 25 s since the robofrog was obscured, females showed a significant preference for the speaker where they could previously observe the robofrog model (binomial test: 18:7; *p* = 0.0361; Fig. 5).

We tested 147 individual frogs across all treatments. Six video recordings were lost or corrupted, so locomotor behaviors were scored on the remaining 141 trials. We categorized each trial into one of two groups: direct paths to the speaker (*n* = 99) or indirect paths (*n* = 42). Females taking an indirect path turned significantly more often (x = 11.26 turns ± 0.97 SE) than females taking a direct path (x = 5.64 turns ± 0.33 SE) (Kruskal-Wallis H = 27.46; *p* = <0.0001; Fig. 6a). Similarly, females taking an indirect path stopped significantly more often (x = 20.40 stops ± 1.67 SE) than females taking a direct path (x = 11.52 stops ± 0.63 SE), (Kruskal-Wallis H = 30.64; *p* = <0.0001; Fig. 6b). In the treatments where there was no silent period, females took a direct path to their speaker of choice significantly more often than an indirect path (binomial test: Robofrog Control, *p* = 0.0004; Blind + Hold, *p* = 0.006; Fig. 6c), with an exception being the Blind Control treatment wherein stimuli were equal (binomial test: *p* = 0.075; Fig. 6c). In treatments with a silent period, females did not take a direct path to their speaker of choice significantly more often than an indirect path (binomial test: Blind + Silence, *p* = 0.922; Blind + Silence + Hold, *p* = 0.091; Fig. 6c).

**Fig. 6.**
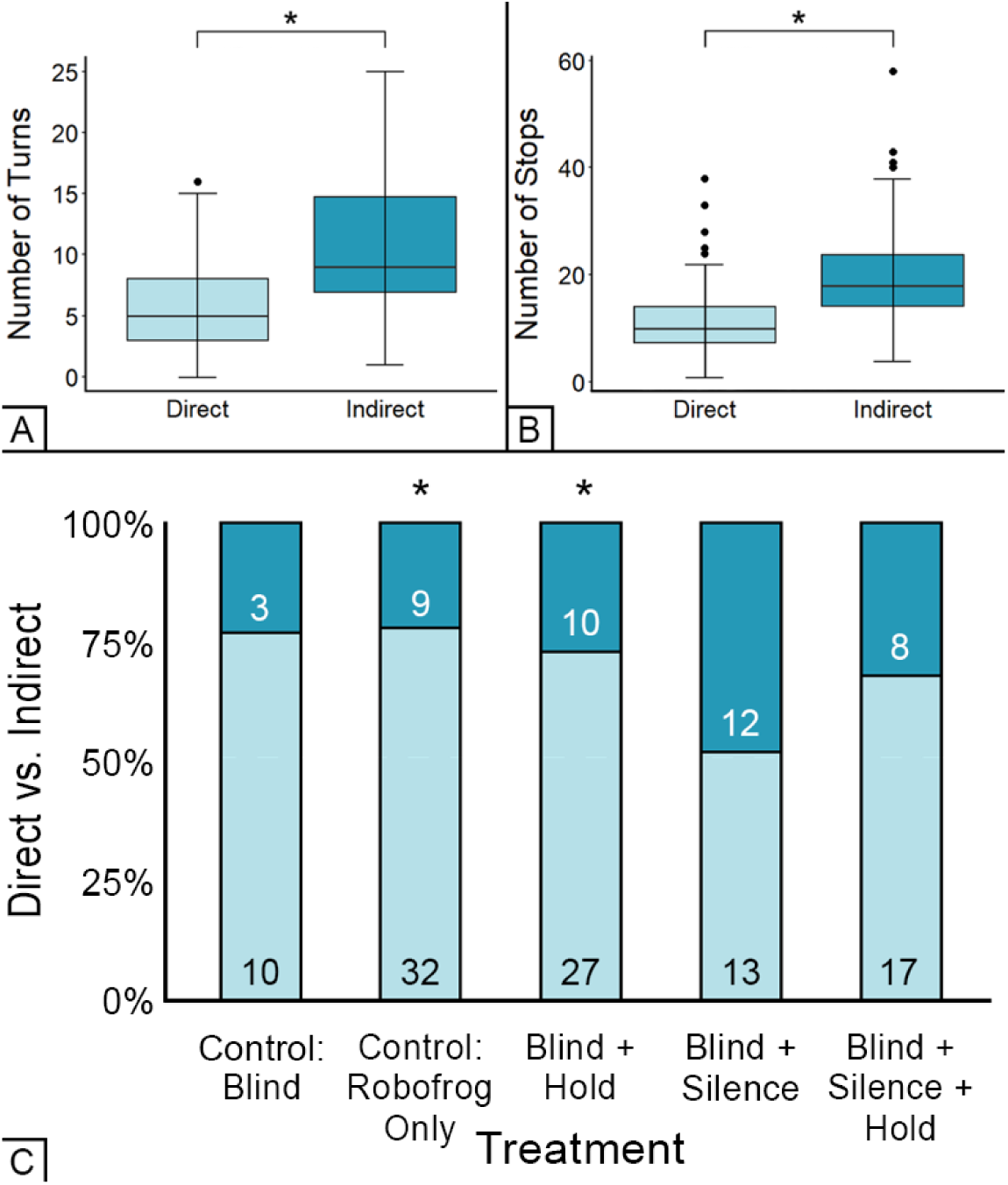
Female locomotor behaviors across treatments. (A) The distribution of turns made by females grouped by whether their path to choice was direct (light blue) or indirect (dark blue). (B) The distribution of stops made by females grouped by whether their path was direct (light blue) or indirect (dark blue). (C) Proportion of females making an indirect/direct choice per treatment. For treatments that did not have a silent period (i.e. Control: Robofrog Only and Blind + Hold), females were significantly more likely to take a direct path (light blue) than an indirect path (dark blue). Numbers inside the bars are the number of individuals that made a direct path or indirect path to choice. Asterisks (*) represent *P* < 0.05.

We also investigated whether the latency of a female to make a choice was affected by treatment type (Fig. 7). All latency data were log-transformed to meet the assumptions of parametric tests. Treatment type did not have a significant effect on choice latency (ANOVA: F_4,136_ = 0.808, *p* = 0.522). Frogs taking a direct path, however, exhibited significantly shorter latencies (x = 46.4 s ± 3.83 SE) than those taking an indirect path (x = 61.45 s ± 7.77 SE) (t_1,139_ = 2.972; *p* = 0.004).

**Fig. 7.**
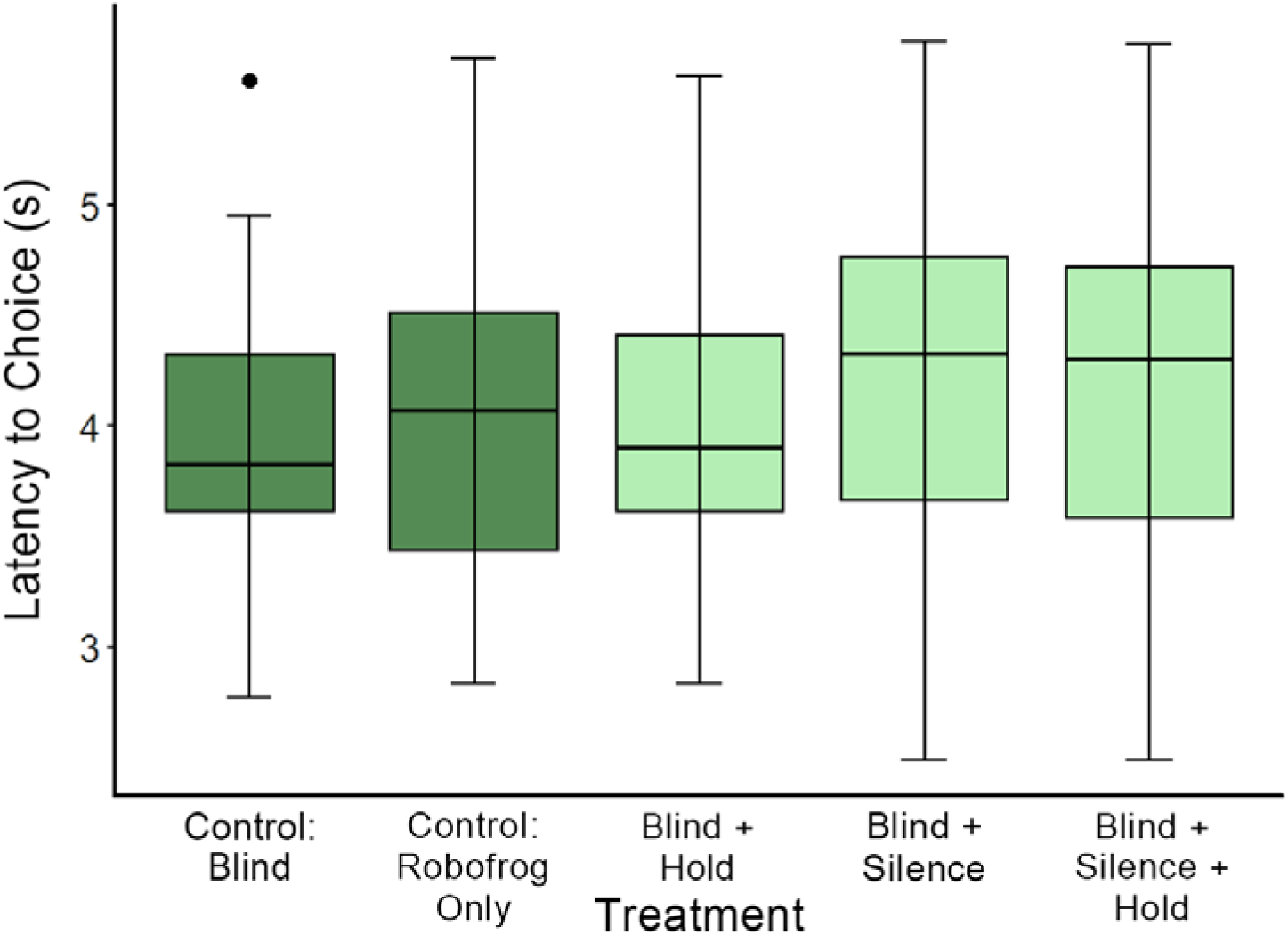
Mean latency to choice across treatments using log-transformed data. The ANOVA did not report a significant difference between any of the treatments’latencies (*p* = 0.964). The dark green boxes (left) represent control treatments.

## DISCUSSION

Because the accessibility of mating displays can vary both temporally and spatially, the ability of females to recall a previously sampled mate is an important, if underappreciated, component of mate choice. While it is to be expected that females within a bonded pair are capable of mate recognition (Miller 1979; Clark et al. 2006; Podraza et al. 2024), we know less about the general capability of females to recall individual males during mate searching. When choosing among multiple potential mates, it would be advantageous to preserve information about individuals so that females can return to certain males and/or have that information to rely on if there is a lapse in accessibility. For some species, this may mean preserving information for an extended period of time (Bensch & Hasselquist 1992; Witte & Massmann 2003), while others may only need to retain this information for less than a minute (Akre & Ryan 2010; Zhu et al. 2021). The ability to recall a single male’s location becomes even more pertinent for females attending a lek as they sample potentially dozens of other males. Amidst a cacophony of rivals, males may increase their chance of being remembered, and therefore chosen, simply by augmenting their display and instantiating memory in sampling females; the noisier, brighter, more flamboyant a male is, the more memorable he may be.

Here, to determine the effect of a multisensory display on the memory of female túngara frogs, we asked if females could remember a “calling male” after the visual component of the male’s display – the dynamically inflating robofrog – was no longer visually accessible. To do this, we developed a pulley system for the lowering of two opaque black cloth blinds that visually obscured both speakers (and the robofrog) from the females’ view. After a presentation period, we gave females either a silent period of 5 s, a holding period of 20 s, or both in sequence. Once the period(s) ended, we released the female from an acoustically and visually transparent funnel and allowed her to make a choice.

Our results provide evidence that the addition of a visual stimulus to a male frog’s courtship call instantiates memory for the caller’s location. Specifically, when females were exposed to an additional minute of stimuli and 5 s of silence before being released, they significantly preferred the speaker that was formerly paired with a robofrog (multimodal speaker) over the bare speaker. However, in the hold treatment (where the audio playback was uninterrupted), females did not exhibit a significant preference for multimodal speaker after last being exposed to the multisensory cues 20 s before release. Because they did seem to remember the location of the cue with exposure to 5 s of silence, we were interested in knowing if this was simply a product of the very short, 5 s delay since females’ last exposure to the robofrog. In the combined Blind + Silence + Hold test, females were still able to remember even after a 25 s delay (5 s of silence plus 20 s delay under the release funnel) and expressed a significant preference for the multimodal speaker.

Here, we point out that the visual component appears to instantiate memory for a caller’s location (a significant proportion of females returned to the speaker where they last saw the robofrog), but in all experimental treatments, females still received auditory information about the location of the speakers. As such, the degree to which females retained spatial memory for a caller’s location is unclear. Burmeister (2022) showed that túngara frogs were capable of learning in a two-choice spatial discrimination task, albeit not as well as poison frogs. Regardless, the females’ choices in our experiment indicate that the visual component influenced their memory such that most females expressed their preference for a multisensory display even when the visual component of that display was removed.

These findings demonstrate a potential mechanism wherein more complex displays may improve a receiver’s ability to remember a signaler, as proposed by Guilford & Dawkins (1991; Rowe 1999). This mechanism does not appear to be species specific, as Zhu et al. (2021) demonstrated a similar effect with a species of treefrog native to Southeast Asia. In that study, researchers were interested in learning how long a multimodal display could instantiate memory, similar to the methods employed by Akre & Ryan (2010) for complex calls in túngara frogs. They found that females were able to recall a caller’s location for 30 s, which was similar to the 45 s of túngara frogs (Akre & Ryan 2010; Zhu et al. 2021). Both of those studies utilized silent periods to titrate the length of time for which the frogs could retain their memory. However, we note that the additional minute of presentation during treatments with a silent period (3 min) versus those without a silent period (2 min) may also play a role in memory instantiation. It is possible that the additional minute of presentation time influenced memory, either alone or in conjunction with the silent period. Although a 3-min “acclimation” period has traditionally been utilized in túngara frog phonotaxis studies (Bosch et al. 2000; Marsh et al. 2000; Baugh & Ryan 2011), future work should aim to disentangle these potential influences. Regardless, our results provide robust evidence that multisensory male stimuli can instantiate memory in females for an individual signaler.

Why a silent period may play a role in memory instantiation could be due, at least in part, to the temporal updating that female túngara frogs appear to exhibit. When male túngara frogs continuously duet with their neighbor (or two duetting speakers in an experiment), females seem to ignore previously available information and make rapid assessments of their options in real-time (Baugh & Ryan 2010a). Thus, females are able to continually modify their choice, but are therefore also sensitive to shifts in the chorus. The sensory scene in a mating chorus is dynamic, with silence itself being a natural part of many chorus soundscapes (Schwartz 1991; Wilson et al. 2014; Greenfield et al. 2016; Coss et al. 2020). Male signalers commonly go silent in response to perceived predation threats, making themselves less conspicuous (Faure & Hoy 2000), but in doing so, they may also be unintentionally activating the female’s ability to remember them. While males are continuously calling, their location is likely discernible to females in at least one modality. When it is already at their disposal, females do not need to retain such spatial information until there is a lapse in the chorus, and so it is likely to be discarded in lieu of new information (Ryan et al. 2009). The temporary cessation of sound, however, seemed to trigger females to remember callers they previously sampled. As previous studies have suggested, it seemed that in the absence of silence (i.e. continual calling), females appeared to continue to update the acoustic signal information in real time and ignore where they had previously seen the calling robofrog. Overall, there was a trend for the non-silent treatment to choose the multimodal speaker, but the lack of silence appears to reduce that response.

As female túngara frogs are making mate choice decisions, we commonly observe them moving and reorienting towards a speaker each time it broadcasts a call. Unsurprisingly, when we evaluated these locomotor behaviors, we found that females that took an indirect path made more stops and turns. These indirect females also displayed longer latencies prior to their choice. More notably, the number of females taking indirect paths was significantly higher in the silent treatments than in those with continued calling. This suggests that the task was cognitively more difficult in the silent treatments and females were relying more on memory to make their choices. If female túngara frogs utilize landmark cues during spatial learning, it is possible that the visual stimulus of an inflating vocal sac, in tandem with calls, acts as a landmark cue for a female to recall a male’s location. Rather than defaulting to a random choice between two unseen males, it seemed that females exposed to a silent period were relying on memory to locate a potential mate whose location had been previously established.

What remains unknown is how the length of the silent period influences memory for caller location. The silent period we employed in this study was 5 s; in nature, males go silent for highly variable periods of time, with mean inter-chorus intervals of 25 s (Akre & Ryan 2010; Dapper et al. 2011). Females may respond strongly to this cue and, as a result, alter their behavior while mate sampling. When danger is sensed, instead of completely “starting over,” females may continue with their decision-making, albeit with more caution (Bernal et al. 2007; Edomwande & Barbosa 2020; Feagles & Höbel 2022).

Previously, a female túngara frog’s memory during mate evaluation had only been tested in one modality: auditory. Akre and Ryan (2010) tested female túngara frogs’ memory for both a simpler call (whine plus one chuck) and a more complex call (whine plus three chucks). They showed that females were only able to remember the location of a speaker when the more complex call (whine plus three chucks) was broadcast. In the present study, a synthetic whine and one-chuck call was used for all acoustic stimuli, and yet, we were still able to induce memory. What both our study and Akre & Ryan (2010) demonstrate is that in order to instantiate memory for signaler location, the display needs to be complex in nature, whether that complexity arises from additional acoustic stimuli (chucks) or from additional modalities (visual). What our findings also highlight is the role that vision plays during mate choice in anurans. It has been proposed that visual stimuli may act to attract the attention of a receiver to that signaler (Ord & Stamps 2008; James et al 2022). In túngara frogs, not only does the visual stimulus attract the receiver’s attention, but it also acts to make the signaler more memorable. Most studies on anuran communication focus on the importance of the acoustic component of the male’s display, but there is increasing evidence that while the acoustic signal is both necessary and sufficient for mate attraction, the visual stimulus of the vocal sac also serves an additional important function (Taylor & Ryan 2013; Starnberger et al. 2014b; Zhu et al. 2022). Females themselves become more visually sensitive when in a reproductive state, which may have evolved to aid them in mate searching and evaluation (Cummings et al. 2008; Leslie et al. 2020). In the absence of a call, the visual component of a túngara frog male’s display does not induce mate searching behavior (Taylor et al. 2011), but here we show that, in tandem with a call, an inflating vocal sac can induce memory for a caller’s location.

Anurans are renowned for their diversity of behavioral and life-history strategies (Wells 2007), so it is not surprising that some species may not rely on visual cues during communication (Li et al 2022; Augusto-Alves et al. 2024). However, we point out that Li et al. (2022) mentioned the use of dim red light when handling frogs in their experiment. Buchanan (1993) demonstrated that dim red light diminishes scotopic visual sensitivity in Cope’s gray treefrogs. Therefore, Li et al. (2022) may have compromised the visual capabilities of the frogs in their experiment. We carefully calibrated our light to fall within the range (both spectrum and intensity) of what túngara frogs experience in nature. Additionally, all frogs remained in complete darkness before and during our experiment, thereby preserving their visual sensitivity. While the anuran vocal sac was traditionally thought to be related only to call production (Bucher et al. 1982; Pauly et al. 2006), several lines of evidence point to the co-option of the vocal sac as a visual component in the communication of a variety of species (Starnberger et al. 2014). First, several species have been shown to preferentially respond to a vocal sac when coupled with a call (Narins et al. 2003; Taylor et al. 2007; Taylor et al. 2008; Gomez et al. 2011; Preininger et al. 2013; Laird et al. 2016; Zhu et al. 2021), and even when not coupled with a call (Hirschmann et al. 2006). Second, the vocal sac varies tremendously in size and color across species (Starnberger et al. 2014; Elias-Costa & Faivovich 2025) and in ways that are not directly linked to sound production. In túngara frogs, the vocal sac is too large to fully inflate unless males are floating in water; this is not true of most frogs (Halfwerk et al. 2017). Despite its large relative size, the túngara vocal sac does not serve as a cavity resonator (Rand & Dudley 1993) and it is too small to effectively transmit the frequencies contained in the whine (Ryan 1985). In addition, frogs similar in size to túngaras, but with smaller vocal sacs (e.g. *Dendropsophus ebrecatus*), produce vocalizations at roughly the same amplitude. We have demonstrated elsewhere that female túngara frogs exhibit a preference for the multisensory display and here again demonstrate a memory function of this display.

Though the focus of this study was not to investigate how long a female túngara frog is able to remember, it is important to note that the length of memory for the location of a calling male in the present study correlates with the previous findings of Akre & Ryan (2010). In that study, the average inter-chorus interval where males would become silent was 25 s. In our study, the longest duration after the multimodal stimulus was last presented was also 25 s, a combination of the previous two memory experiments’ treatment periods. When presented with a complex acoustic-only stimulus, female túngara frogs were able to remember its location for up to 45 s (Akre & Ryan 2010); how that duration compares to a multimodal stimulus remains unknown. In a treefrog species, Zhu et al. (2021) demonstrated that multimodal stimuli generate a longer working memory. Zhu et al. (2022) also showed that the presence of an inflating vocal sac plus noise, but not either stimulus alone, restores salience to calls with missing syllables. Future studies are needed to further compare the memory capabilities between unimodal and multimodal stimuli and across additional taxa.

Our understanding of the active role that females play in sexual selection has exponentially increased since the time of Darwin (Rosenthal & Ryan 2022). There are various hypotheses to explain why females generally prefer more complex stimuli; the receiver psychology hypothesis has provided a simple and powerful explanation for the mechanism of female choice for multimodal displays (Rowe 1999). Interestingly, despite being proposed more than two decades ago, the role of multimodal cues instantiating memory in receivers has received relatively little attention. The evolutionary currency is fitness; thus, it follows that the more complex a display the more memorable it will be for the receiver and therefore be under positive selection. In the present study, while we found that multimodality was important for memorability, in order for memory to be induced, an additional minute of presentation along with a potential disturbance/predation risk (silent period) was necessary for memory instantiation. In a lekking scenario, females continue to make a choice despite this risk, likely so as to not lose their reproductive investment (Bernal et al. 2007). We know that females dynamically assess males (Baugh & Ryan 2010a,b) and will circle back to them once a “final” choice has been made (Dale & Slavsgold 1996). An association in a female’s brain between one particular male and his location must therefore exist. Multimodal stimuli have been shown to improve both memory and learning outside of the context of mate choice (foraging: Leonard et al. 2011; Gil-Guevara et al. 2022; aposematic signals: Speed 2000; Rowe 2002; Siddall and Marples 2008; and song-learning: Hultsch et al. 1999; Varkevisser et al. 2022). By utilizing a multisensory display, signalers may therefore increase their chances of being remembered. Thus, how a signaler is chosen may be determined not by how well they advertise quality, but simply by how memorable they are during fluctuations in a dynamic signaling environment. Here, we used túngara frogs to better understand how the use of multiple sensory modalities may influence a female’s memory and subsequent mate choice. Our results reflect the complex decisions females must make during mate evaluation, especially when multiple communication modalities must be integrated and later remembered. A receiver’s cognitive ability to adapt to rapidly shifting environments and still retain the ability to weigh attractiveness is both impressive and meaningful. The choice a female makes has direct evolutionary consequences, and furthering our understanding of the process of decision-making is critical for understanding the evolution of diversity.

## ACKNOWLEDGEMENTS

We thank Jorge López and Leslie Barría for their contribution with both field work and data collection. We also want to thank Gregg Cohen and Luke Larter for their support during field seasons. Paul Clements supported our team by designing the robotic units and Christopher Maness aided with graphic design. Nana Osei-Wusu served as the blind observer for scoring videos. Golden Frog Scuba provided compressed air cylinders that powered the pneumatic robofrog control units. The Smithsonian Tropical Research Institute provided logistical support and laboratory space. We are grateful to the Republic of Panamá for hosting our team.

## Notes

https://doi.org/10.60635/C3GP6F

### Competing Interest Statement

The authors have declared no competing interest.

### Summary of Updates

This version of the manuscript has been revised to update the following: Acknowledgements added.

### Funding

- National Science Foundation: IOS-1914646

